# Implications of data-driven analyses for personalized therapy in psychosis: a systematic review of cluster- and trajectory-based modelling studies

**DOI:** 10.1101/599498

**Authors:** Tesfa Dejenie Habtewold, Lyan H. Rodijk, Edith J. Liemburg, Grigory Sidorenkov, H. Marike Boezen, Richard Bruggeman, Behrooz Z. Alizadeh

**Author notes:** =Corresponding author Behrooz Z. Alizadeh MD, MSc, PhD, University of Groningen, University Medical Center Groningen, Department of Epidemiology, Hanzeplein 1, 9713 GZ, Groningen, The Netherlands.

## Abstract

**Introduction:** To tackle the phenotypic heterogeneity of schizophrenia, data-driven methods are often applied to identify subtypes of its (sub)clinical symptoms though there is no systematic review.

**Aims:** To summarize the evidence from cluster- and trajectory-based studies of positive, negative and cognitive symptoms in patients with schizophrenia spectrum disorders, their siblings and healthy people. Additionally, we aimed to highlight knowledge gaps and point out future directions to optimize the translatability of cluster- and trajectory-based studies.

**Methods:** A systematic review was performed through searching PsycINFO, PubMed, PsycTESTS, PsycARTICLES, SCOPUS, EMBASE, and Web of Science electronic databases. Both cross-sectional and longitudinal studies published from 2008 to 2019, which reported at least two statistically derived clusters or trajectories were included. Two reviewers independently screened and extracted the data.

**Results:** Of 2,285 studies retrieved, 50 studies (17 longitudinal and 33 cross-sectional) conducted in 30 countries were selected for review. Longitudinal studies discovered two to five trajectories of positive and negative symptoms in patient, and four to five trajectories of cognitive deficits in patient and sibling. In cross-sectional studies, three clusters of positive and negative symptoms in patient, four clusters of positive and negative schizotypy in sibling, and three to five clusters of cognitive deficits in patient and sibling were identified. These studies also reported multidimensional predictors of clusters and trajectories.

**Conclusions:** Our findings indicate that (sub)clinical symptoms of schizophrenia are more heterogeneous than currently recognized. Identified clusters and trajectories can be used as a basis for personalized psychiatry.

## Introduction

In psychiatry, one of the major challenges for tailoring individualized therapies are phenotypic heterogeneity of disorders and its overlapping symptoms that may presumably share some fundamental biologic underpinnings.^1^ In schizophrenia, a complex psychotic disorder that affects individuals and families, the phenotypic expression and course of disease are variable.^2^ The prevalence of schizophrenia is 4.6 per 1.000 individuals with a lifetime morbidity risk of 0.7%.^3^ The twin- and SNP-based heritability estimate of schizophrenia was 80%^4^ and 30%^5^, respectively. The clinical symptoms of schizophrenia are positive symptoms (hallucinations and delusions), negative symptoms (emotional expressive deficit, social amotivation, social withdrawal and difficulty in experiencing pleasure) and cognitive deficits (selective or global).^6^ These symptoms are assessed by standard psychometric tools, which rate symptoms in quantitative scales.^7–12^ The prevalence of negative symptoms is 50-90% in first-episode psychosis and persists in 20-40% of patients with schizophrenia.^13–15^ Cognitive deficits affects 75-80% of patients with schizophrenia.^16^ The most common deficits occur in executive function, processing speed, memory (e.g. episodic, verbal and working), attention, verbal fluency, problem-solving and social cognition.^17–25^ Thus far, patients harbor a wide range of subjectively defined symptoms and phenotypes, which together yields instinctively to heterogeneous groups of people who are collectively diagnosed as schizophrenia. Subclinical symptoms are also evident in siblings of patients with schizophrenia spectrum disorders and healthy general population. ^26–28^

### Heterogeneity in schizophrenia

Despite a century of efforts, understanding the heterogeneity in presentation and course of schizophrenia has been unsuccessful due to the subjective measurement of its clinical symptoms, variation in response to treatment, lack of valid, stable, and meaningful subphenotyping methods, and limited understanding of the disease mechanism.^29–31^ Heterogeneity in clinical outcomes can be manifested within patients and between groups of patients, within subjects over time, and within and between diseases subphenotypes, and caused by several intrinsic and extrinsic factors.^30, 32^ Identification of meaningful homogeneous subgroups of the population based on clinical features or endophenotypes (e.g. neuropsychological markers, neural substrates, and neurological soft signs) requires the use of both supervised and unsupervised analyses. Distinguishing heterogeneous patients to more homogeneous subgroups is expedient not only to unveil common etiologies, but also at practical level to examine the patterns of clinical symptoms, understand the inherent course of the disease, predict treatment response and develop new treatment strategies specific to that subgroup to improve recovery and functional outcomes (Figure 1).^29, 30, 33, 34^

**Figure 1:**
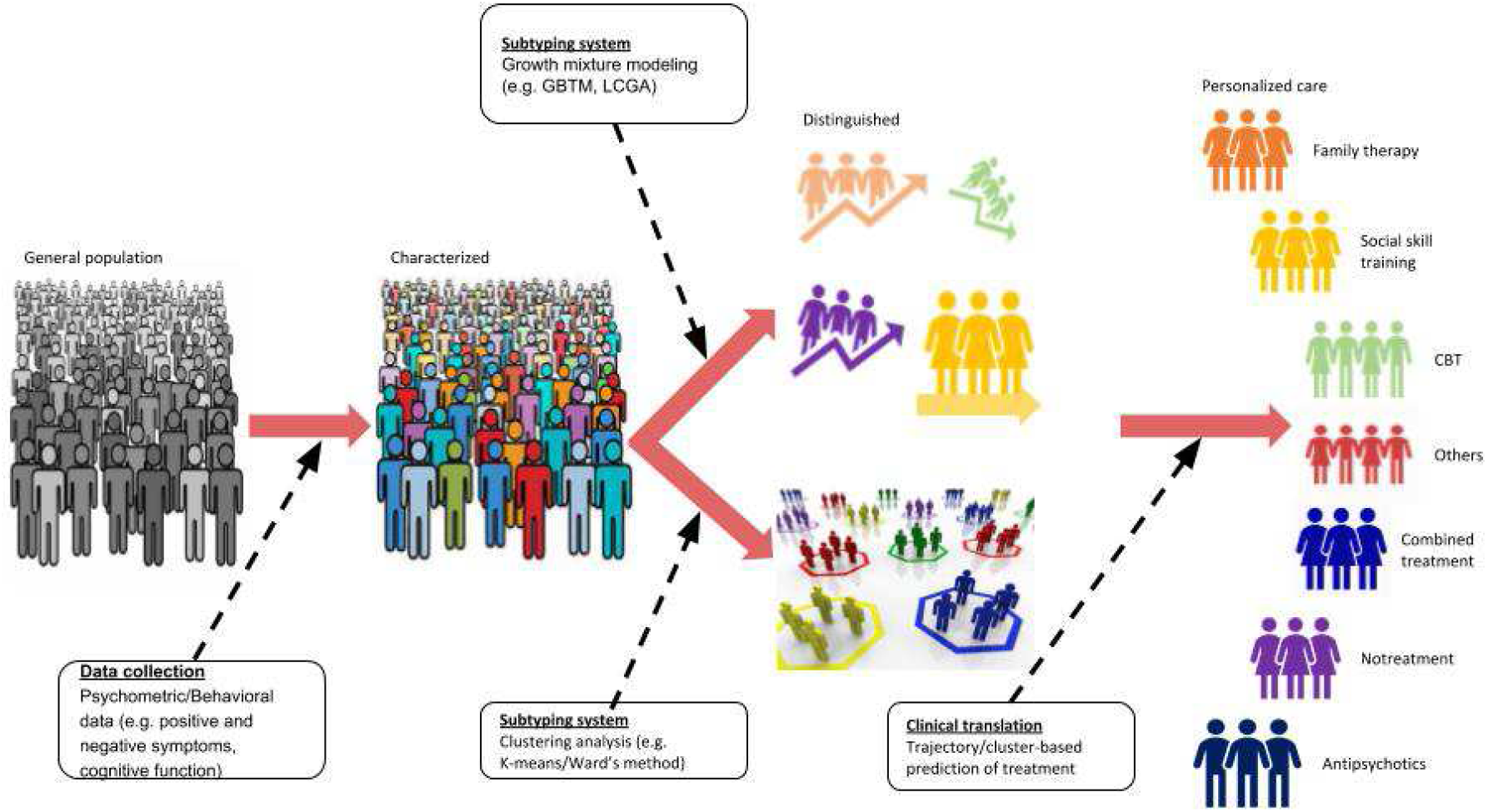
Precision in psychiatric care through measurement, characterization and subtyping.

### Tackling heterogeneity in schizophrenia

For tackling heterogeneity, in the past decade, numerous efforts have been undertaken by carefully designing studies and developing statistical models implemented in various programming language and software.^30^ As a result, clusters or trajectories of clinical symptoms have been estimated using latent class cluster analysis and growth mixture models respectively.^29, 35, 36^ A trajectory or cluster is a group of individuals that has a homogenous symptom profile within that group and a significantly dissimilar profile from other groups.^33^ Statistical methods can identify subgroups and describe within and between-variation that help clinicians and statisticians to explore the relationship of schizophrenia with various clinical and functional outcomes, treatment response, and neuropathological change. Dichotomization of clinical outcomes, such as recovered or not, and symptom remission or not is also a common practice within schizophrenia research.^33^ However, dichotomization may lead to the loss of information, inefficient analysis of continuous data and difficulties in the translation of results to clinically meaningful evidence.^33^ Moreover, subtyping using imaging, biological and symptom data is a recognizable method.^35^

Cluster- and trajectory-based studies of clinical symptoms of schizophrenia show inconsistent findings and have several limitations. Possible reasons of inconsistencies are the heterogeneity of study population, high symptomatic variability between patients and within patients over time, use of various assessment tools, use of different clustering algorithms, and use of different scoring and standardization techniques.^13, 18, 37^ The major limitations are small sample size, short duration of follow-up, and limited used of data from healthy siblings and/or controls.^37^ All these factors blur our understanding of the heterogeneity of the course of schizophrenia. Several reviews have been conducted on cognitive dysfunction^16, 38–47^, negative symptoms^15, 48, 49^ and positive symptoms.^50^ However, these proceedings have largely focused on the traditional approach in determining average change in the course of symptoms over time, and variation between subjects (patient vs sibling, sibling vs control, patient vs control) and diagnosis. They are also based on correlation analysis, which is believed not to be a strong measure of association between predictors and outcomes. In addition, none of these reviews fully addressed symptomatic clusters and trajectories in patients with schizophrenia spectrum disorders, their siblings and healthy controls. Therefore, there is a pressing need to synthesize the contemporary evidence, evaluate the extent and origin of heterogeneity, and to inform personalized and preventive strategies for clinical practice.In this systematic review, we summarized the contemporary evidence from cluster- and trajectory-based studies of positive and negative symptoms/schizotypy, and cognitive deficits in patients with schizophrenia spectrum disorders, their siblings and healthy people. Additionally, we explored the methodological approaches applied to distinguish homogeneous subgroups. We further highlighted current knowledge gaps and point out future directions to optimize the translatability of cluster- and trajectory-based studies within outlooks of personalized approach.

## Methods

### Registration and reporting

This systematic review was conducted and reported based on a registered protocol^51^ and the Preferred Reporting Items for Systematic Review and Meta-Analysis (PRISMA) statement guideline (Supplementary file 1) respectively.^52, 53^ The screening and selection process of the reviewed articles are further illustrated using a PRISMA flow diagram.

### Databases and search terms

A systematic search of PubMed, PsycINFO, PsycTESTS, PsycARTICLES, SCOPUS, EMBASE and Web of Science electronic databases was performed. A comprehensive search strategy was developed for PubMed and adapted for each database in consultation with a medical information specialist (Supplementary file 1). The following search terms were used in their singular or plural form in their title, abstract, keywords and text: “schizophrenia”, “psychosis”, “non-affective psychosis”, “cognitive deficit”, “cognitive dysfunction”, “cognitive alteration”, “negative symptoms”, “deficit syndrome”, “positive symptoms”, “psychopathology”, “cognit*”, “neuropsycholog*”, “neurocognition”, “longitudinal”, “follow-up”, “course”, “heterogeneity”, “endophenotype”, “profile”, “cluster analysis”, “siblings”, “healthy controls”, “latent class analyses”, “Symptom trajectories”, “traject*”, “group modelling” and “trajectory”. Cross-references of included articles and grey literature were also hand-searched. Furthermore, we searched the table of contents of the journals of Schizophrenia Research, Schizophrenia Bulletin, Acta Psychiatrica Scandinavica and British Journal of Psychiatry to explore relevant studies. The freezing date for final search was August 2019. In this review, we use ‘trajectory’ for groups identified by longitudinal studies and ‘cluster’ for groups identified by cross-sectional studies.

### Inclusion and exclusion criteria

Studies meeting the following criteria were included: (1) cross-sectional and longitudinal studies; (2) studies that reported at least two clusters or trajectory groups of individuals using a statistical method based on distinct positive symptom, negative symptom, and neurocognitive or social cognitive impairment dimensions or a combination of these symptom dimensions; (3) studies conducted in patients with schizophrenia-spectrum disorders, and/or their unaffected siblings, and/or healthy individuals irrespective of any clinical (e.g. medication status, severity of illness) and sociodemographic characteristics; and (4) studies published in English from 2008 to 2019. The publication year was limited to the last decade to capture the latest available evidence, which are likely to provide statistically powerful precise estimates and successful subtyping of schizophrenia symptoms due to the increased number of large cohorts. In order to maximize the number of searched articles, the follow-up period in longitudinal studies was not restricted. Trajectory studies based on analyses of the mean level of change for the entire sample were excluded because they did capture individuals’ patterns of change over time and treat between-subject variation as error, so that the actual heterogeneity of groups cannot be revealed.^54^ In addition, studies based on the non-statistical methods of clustering (e.g. family-based clustering) were excluded. Review papers, commentaries, duplicate studies, editorials, and qualitative studies were excluded as well. Furthermore, we excluded studies in which the trajectory groups or clusters were generated based on scores constructed using a combination of schizophrenia symptoms and other unspecified psychotic symptoms.

### Data retrieval and synthesis

Studies retrieved from all databases were exported to RefWorks version 2.0 for Windows web-based citation manager. Close and exact duplicates were deleted. All independent studies were exported to a Microsoft Excel spreadsheet to screen for further inclusion criteria. Authors TD and LR independently screened the titles and abstracts. The two reviewers had substantial agreement, as shown by a Kappa coefficient of 0.62. Inconsistent decisions on title and abstract inclusion were discussed with corresponding author BZA. Finally, full-text was reviewed, and the following data were independently extracted by TD and LR: first author name, publication year, country, cohort/research center, study population, sample size, symptom dimension(s), assessment tool, study design, duration of follow-up for longitudinal studies, frequency of assessment, method of calculating composite score, method of clustering/trajectory analysis, number of identified clusters or trajectory groups and significant predictors of clusters and trajectories.^55^ The corresponding author was contacted by email if full-text of included article was not accessible. If the cohort or research center was not clearly reported, we extracted the institutional affiliation of the first or corresponding author.

## Results

### Search results

In total, 2,262 studies were identified through database searching and an additional 23 studies through manual searching of cross-references and tables of content of relevant journals. After removing duplicate articles and applying the inclusion and exclusion criteria, titles and abstracts of 1,294 articles were screened, resulting in the exclusion of 1,236 articles. In total, 58 articles were selected for full-text review, and eight articles^56–63^ were excluded due to unclear outcome, mixed diagnosis of the study population, use of non-statistical method of clustering or clustering based on different phenotypes of schizophrenia. Finally, data were extracted from 50 cluster- and trajectory-based studies. The PRISMA flow diagram of screening and the selection process is shown in Figure 2.

**Figure 2:**
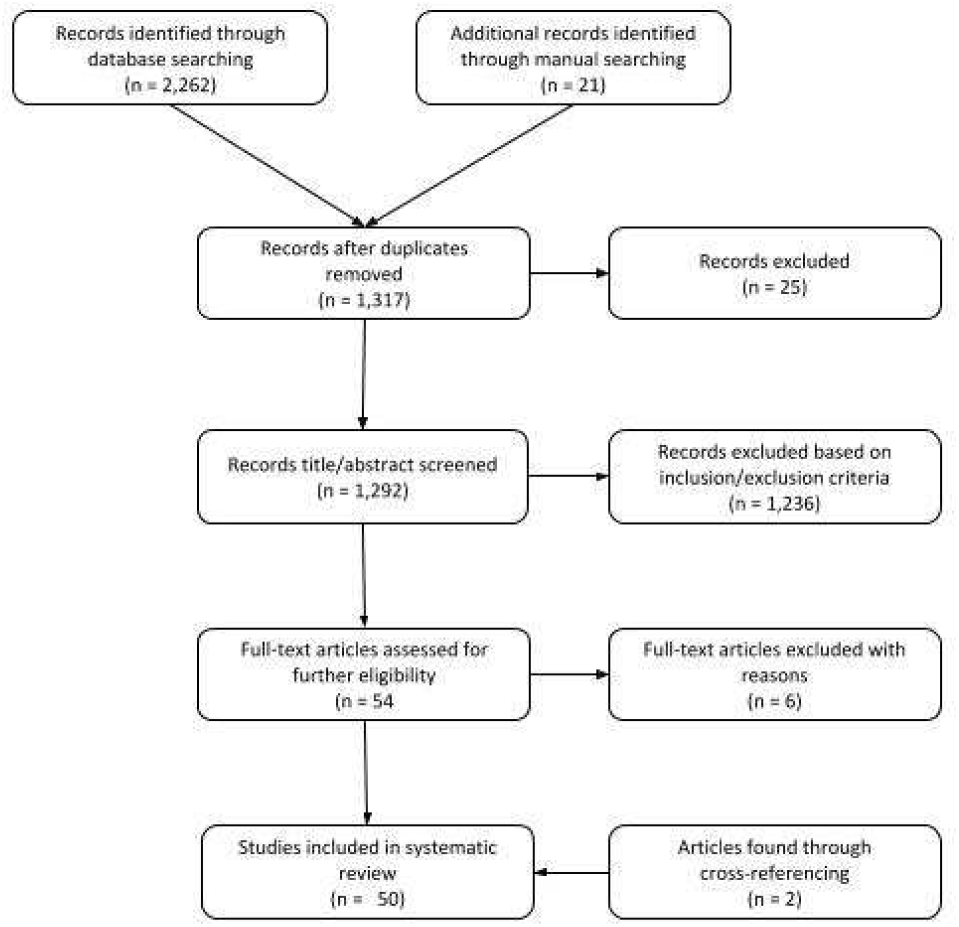
PRISMA flow diagram illustrating the screening and selection of literatures.

### Overview of included studies

The included 50 studies were conducted globally in 30 countries (16 studies in the USA) and published over a decade from 2009 to 2019. Of these, 17 studies were longitudinal that involved 11,475 patients, 1,059 siblings and 2,194 controls/general population, whereas 33 studies were cross-sectional that involved 5,598 patients, 7,423 siblings, and 2,482 controls. Only one longitudinal study^64^ and three cross-sectional studies^65–67^ examined symptomatic subtypes among siblings. Most of the longitudinal studies examined trajectories of positive and negative symptoms, whereas most cross-sectional studies explored clusters based on cognitive function. A minimum of two and maximum of five schizophrenia symptoms subtypes were discovered.

### Symptomatic trajectories

Of the total of 17 longitudinal studies (Table 1), conducted in more than eight countries, 11 studies^31, 33, 34, 36, 68–74^ investigated the trajectory of both positive and negative positive symptoms in patients, three studies^75–77^ the trajectory of only negative symptoms in patients, one study^78^ the trajectory of schizotypy, and two studies^30, 64^ examined the trajectory of neurocognitive impairment in patients and siblings. The duration of follow-up ranged from six weeks to 10 years and included all population age groups. The sample size ranged from 138 to 1,990 subjects, though variation observed between symptom dimension. One study^64^ investigated the association between patients’ and siblings’ cognitive trajectories, whereas another study^74^ examined the association between positive and negative symptom trajectories in patients. Additionally, five studies reported the influence of trajectories on long-term social, occupational and global functioning, and health-related or general quality of life.^34, 73, 75–77^

**Table 1:**
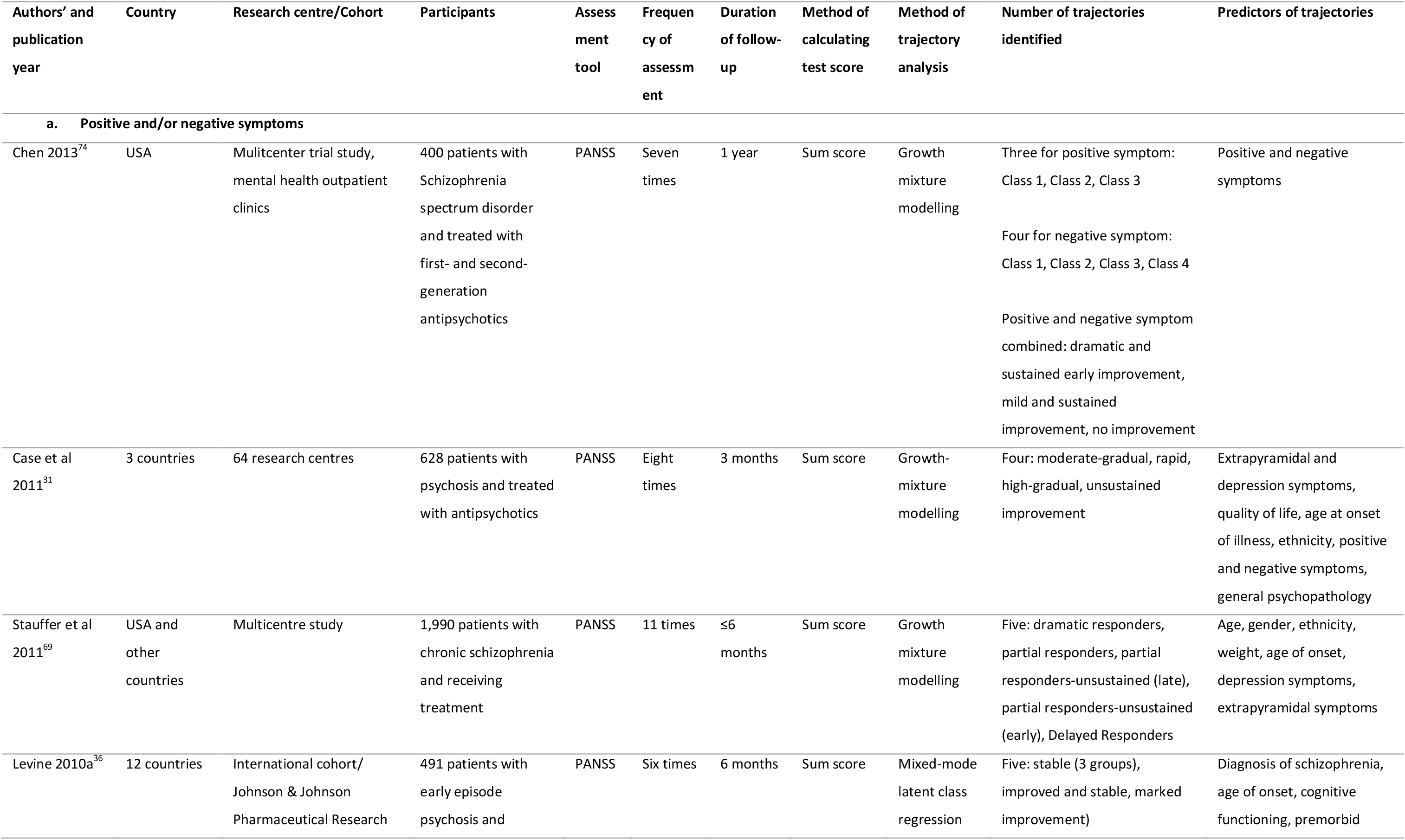
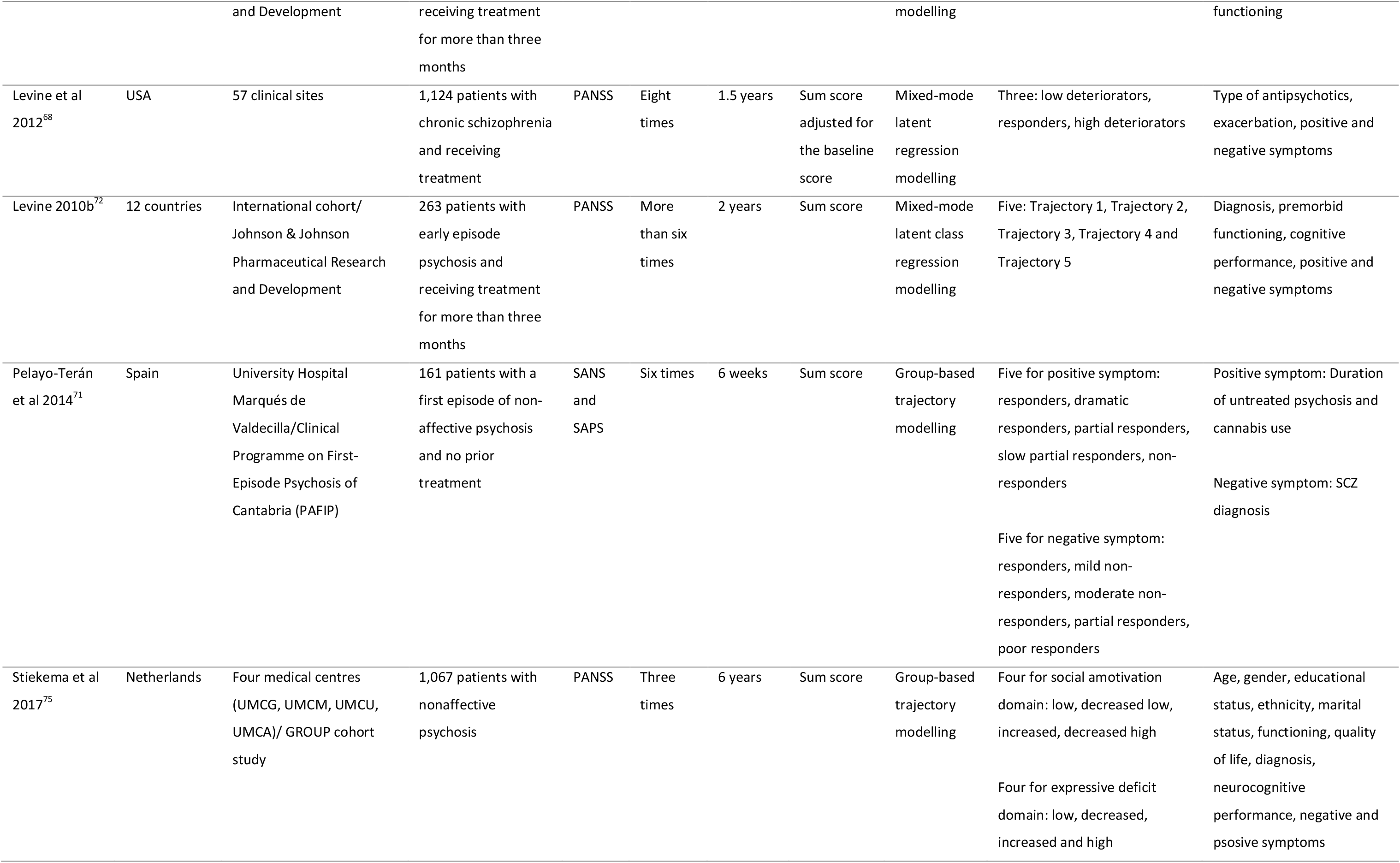
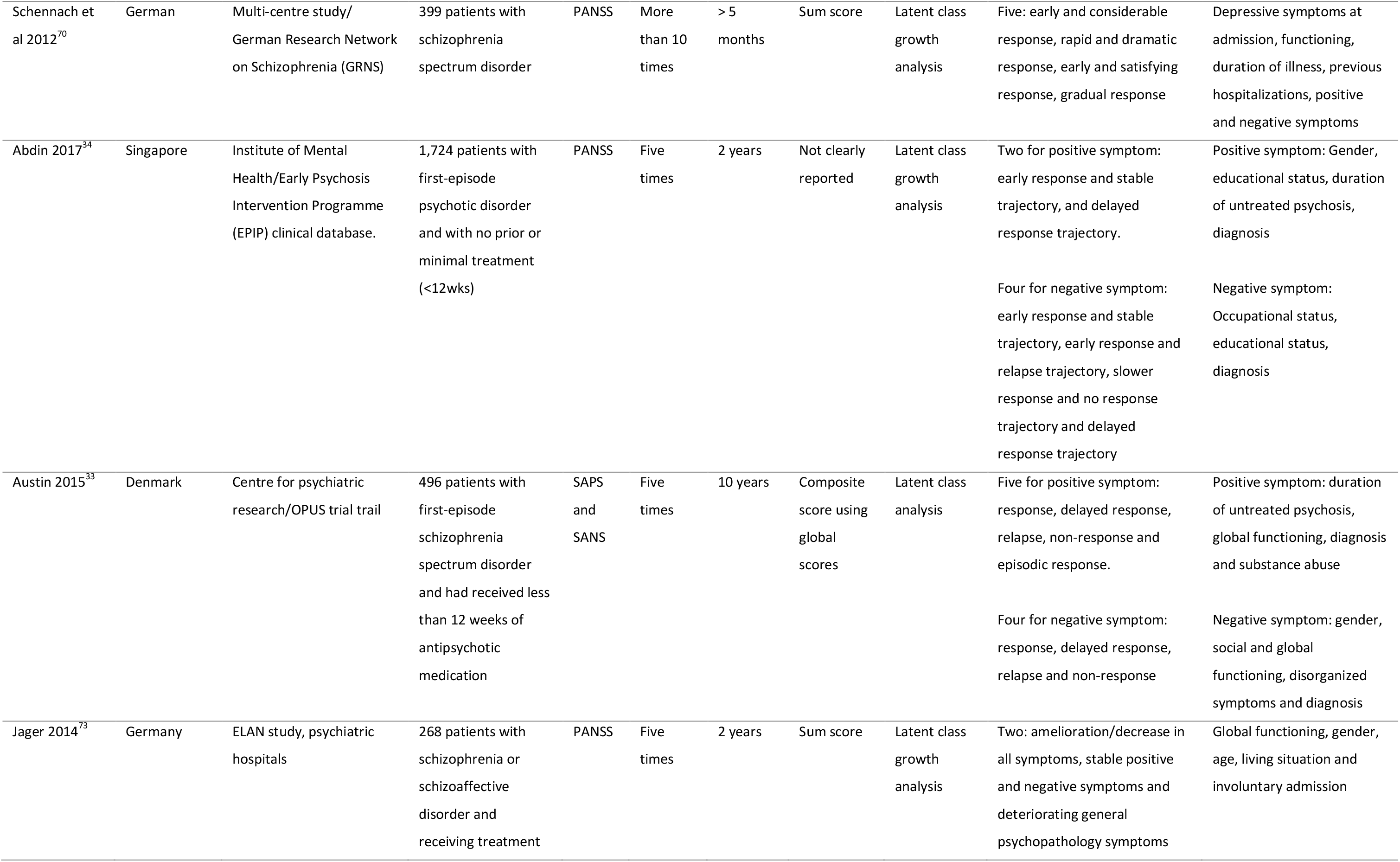
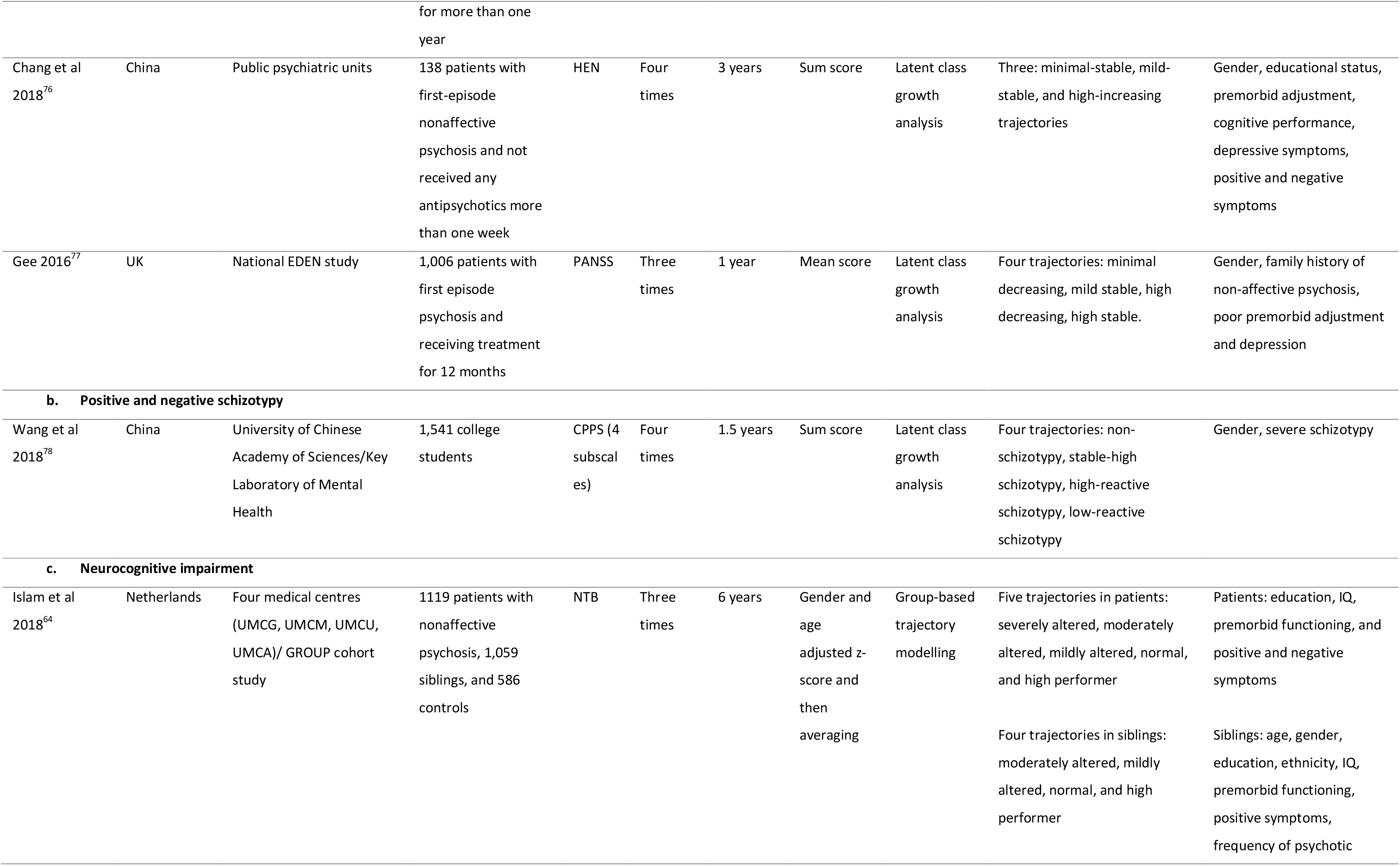
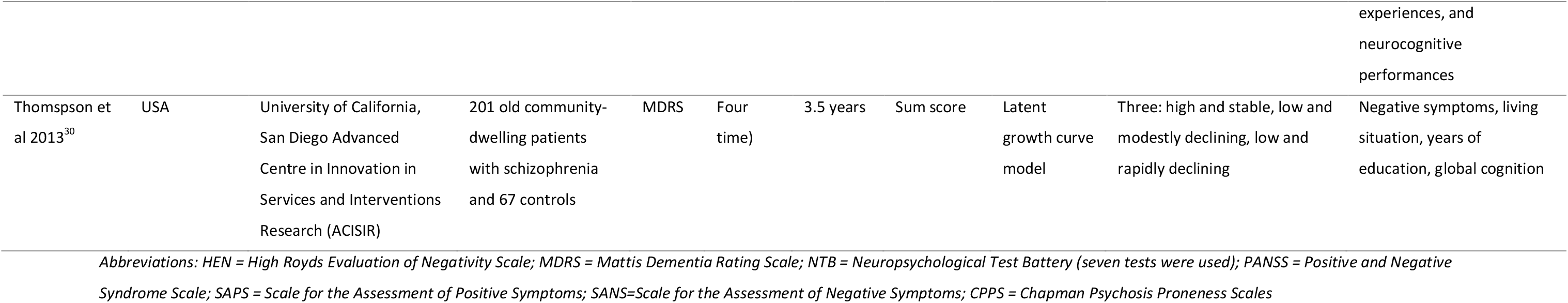
Detailed characteristics of longitudinal studies (n = 17).

Even though all studies had similar aims, they used slightly different methods of trajectory analysis, such as growth mixture modelling (GMM)^31, 69, 74^, latent class growth analysis (LCGA)^30, 33, 34, 70, 73, 76, 77^, mixed mode latent class regression modelling^36, 68, 72^ and group-based trajectory modelling (GBTM).^64, 71, 75^ Akaike’s Information Criterion (AIC), Bayesian information criterion (BIC), logged Bayes factor, sample-size-adjusted BIC (aBIC), bootstrap likelihood ratio test [BLRT], Lo–Mendell–Rubin Likelihood Ratio Test (LMR-LRT) and entropy were reported model selection indices. Of these indices, Bayesian information criterion (BIC) was reported by all studies except for one study^30^ that reported deviance information criterion (DIC).

As shown in Table 1, five studies^33, 36, 69, 71, 72^ discovered five trajectories, three studies^31, 68, 74^ identified three trajectories, and two studies^34, 73^ found two trajectories of positive symptoms. Similarly for the negative symptom dimension, four studies^36, 69, 71, 72^ discovered five trajectories, five studies^31, 33, 34, 74, 77^ reported four trajectories, one study^76^ depicted three trajectories and one study^73^ found two trajectories. In addition, a study^75^ from our research group identified four trajectories of negative symptom subdomains of social amotivation and expressive deficits. Combining both positive and negative symptom dimensions, three studies^36, 70, 72^ discovered five trajectories, one study^31^ found four trajectories and one study^74^ identified three trajectories. One study^78^ identified four trajectories of positive and negative schizotypy in college students without psychosis. With regard to cognitive deficits, a six year longitudinal study^64^ from our research group discovered five trajectories of cognitive impairment in patients and four trajectories in healthy siblings. Another study^30^ reported three trajectories of global cognitive function combining patients and controls together. Overall, these studies characterized trajectories as progressive deterioration, relapsing, progressive amelioration and stable.

### Symptomatic clusters

Of the 49 included studies, 33 studies were cross-sectional conducted in 14 countries (Table 2). The total sample size per study ranged from 62 to 6,600 individuals irrespective of participants’ diagnostic status. Among 32 studies, 21 studies^32, 37, 65, 66, 79–96^ reported clusters in patients and one study^66^ in unaffected siblings based on neurocognitive and/or social cognitive function. In addition, two studies were conducted on negative symptoms^29, 97^, one study on positive symptom^98^, three studies on positive and negative symptoms^35, 99, 100^, and three studies on positive and negative schizotypy.^67, 101, 102^

**Table 2:**
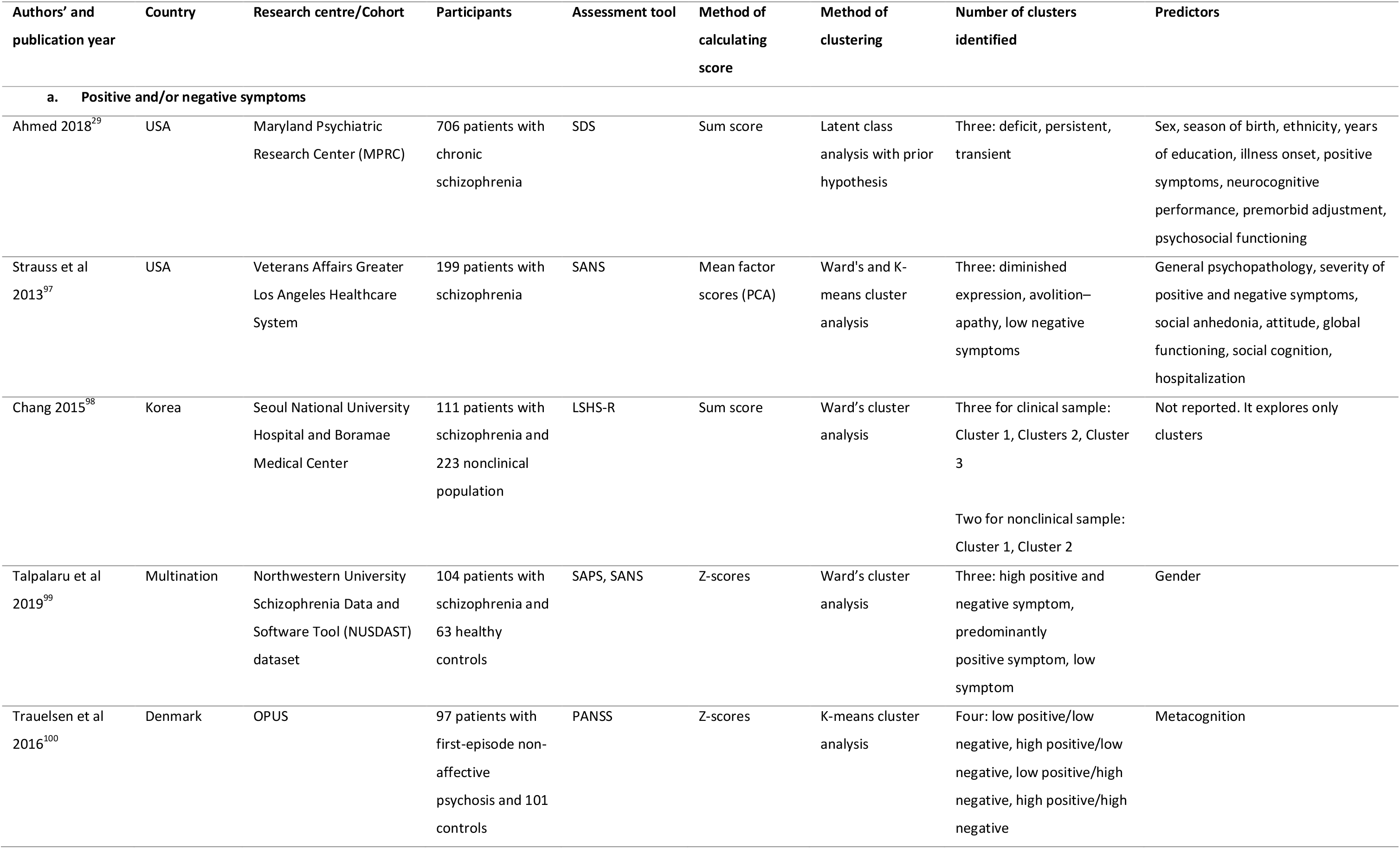
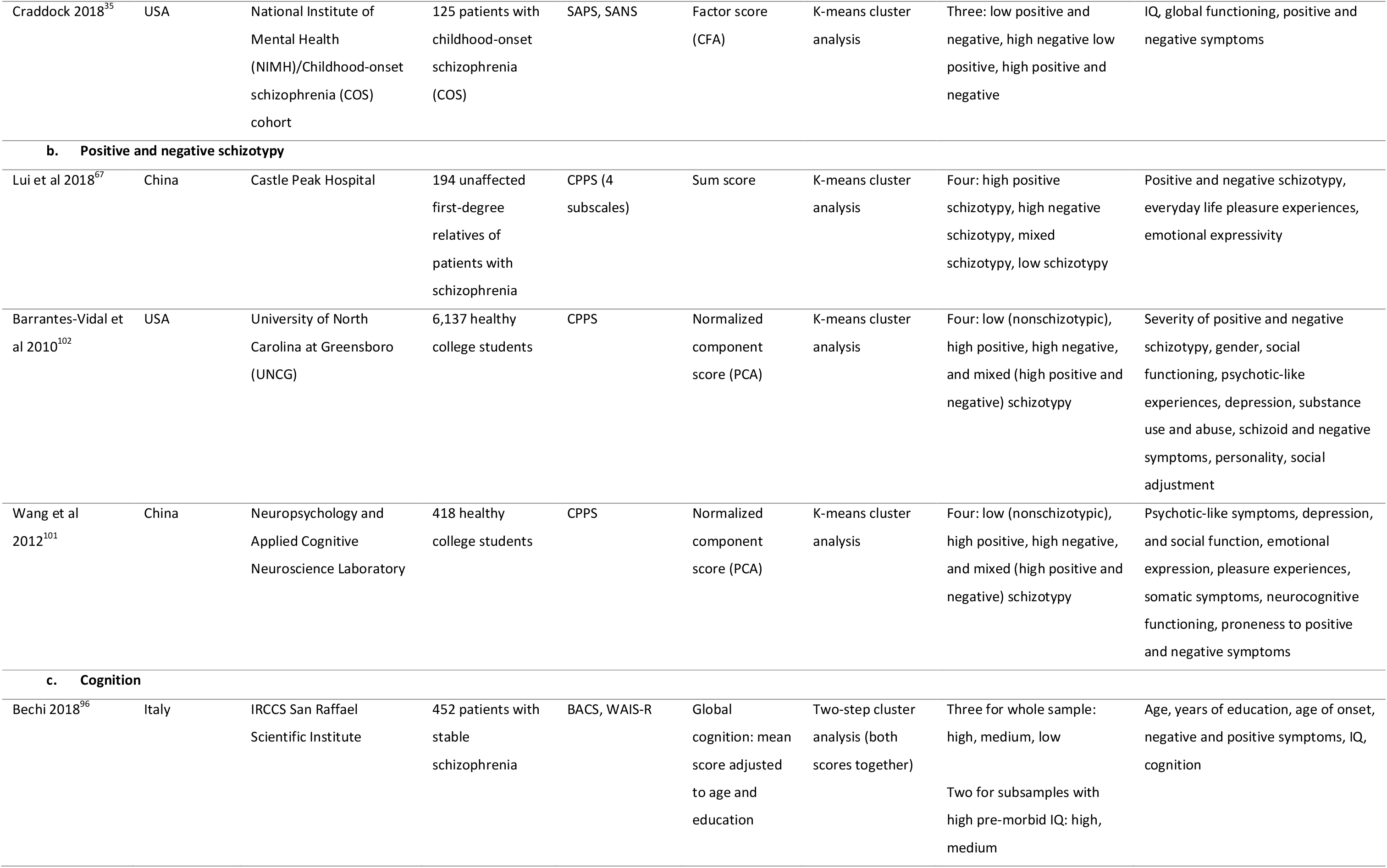
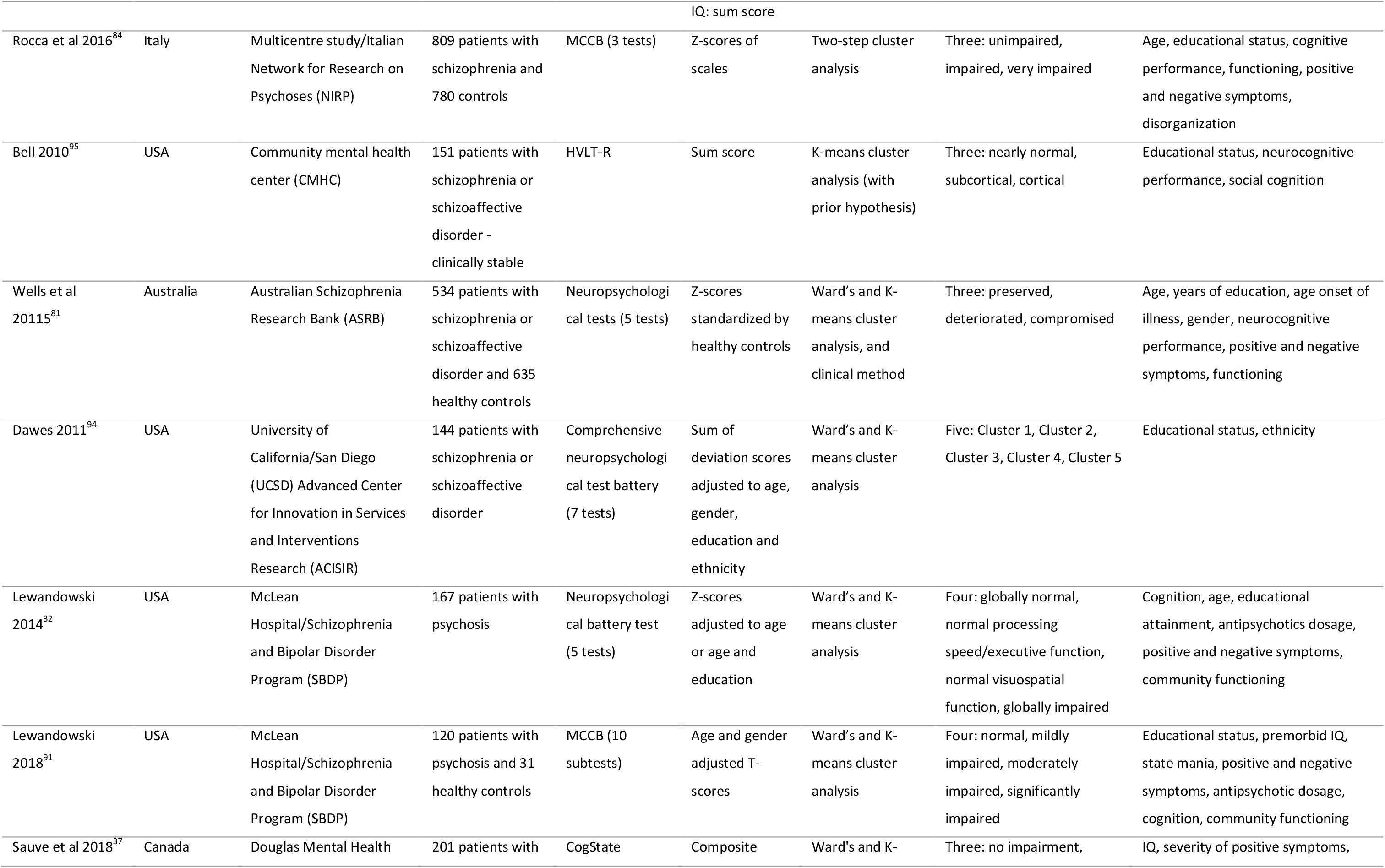
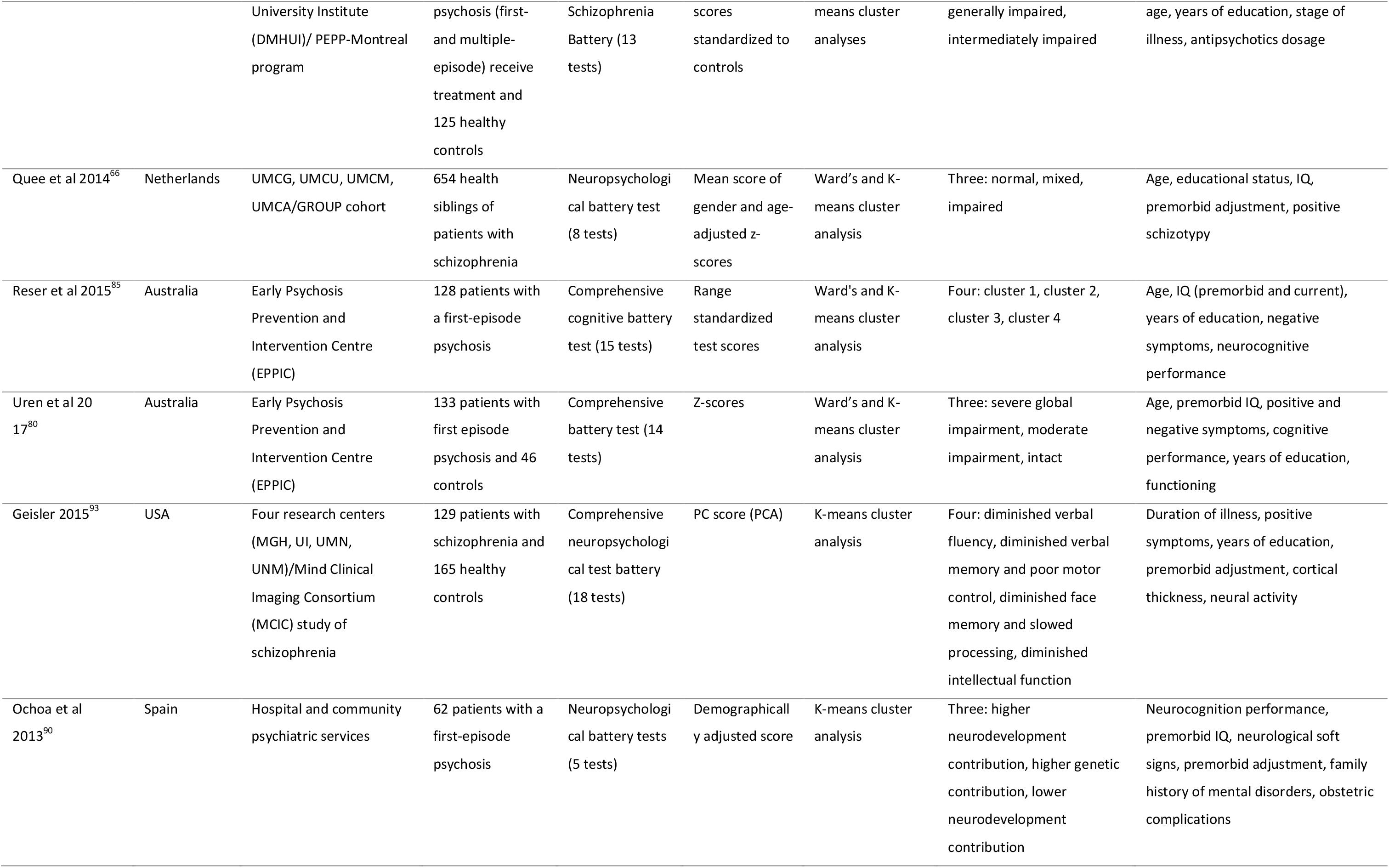
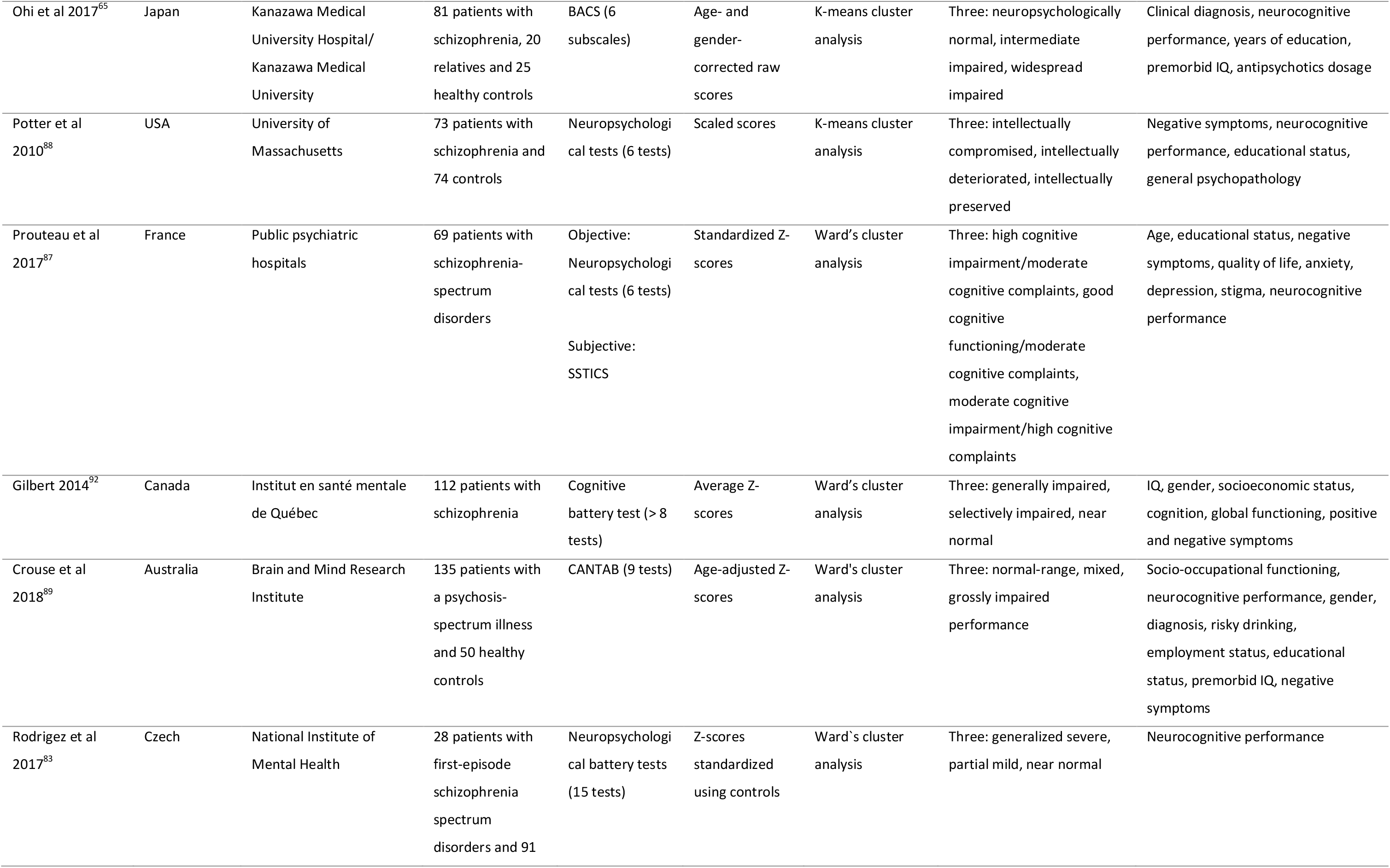
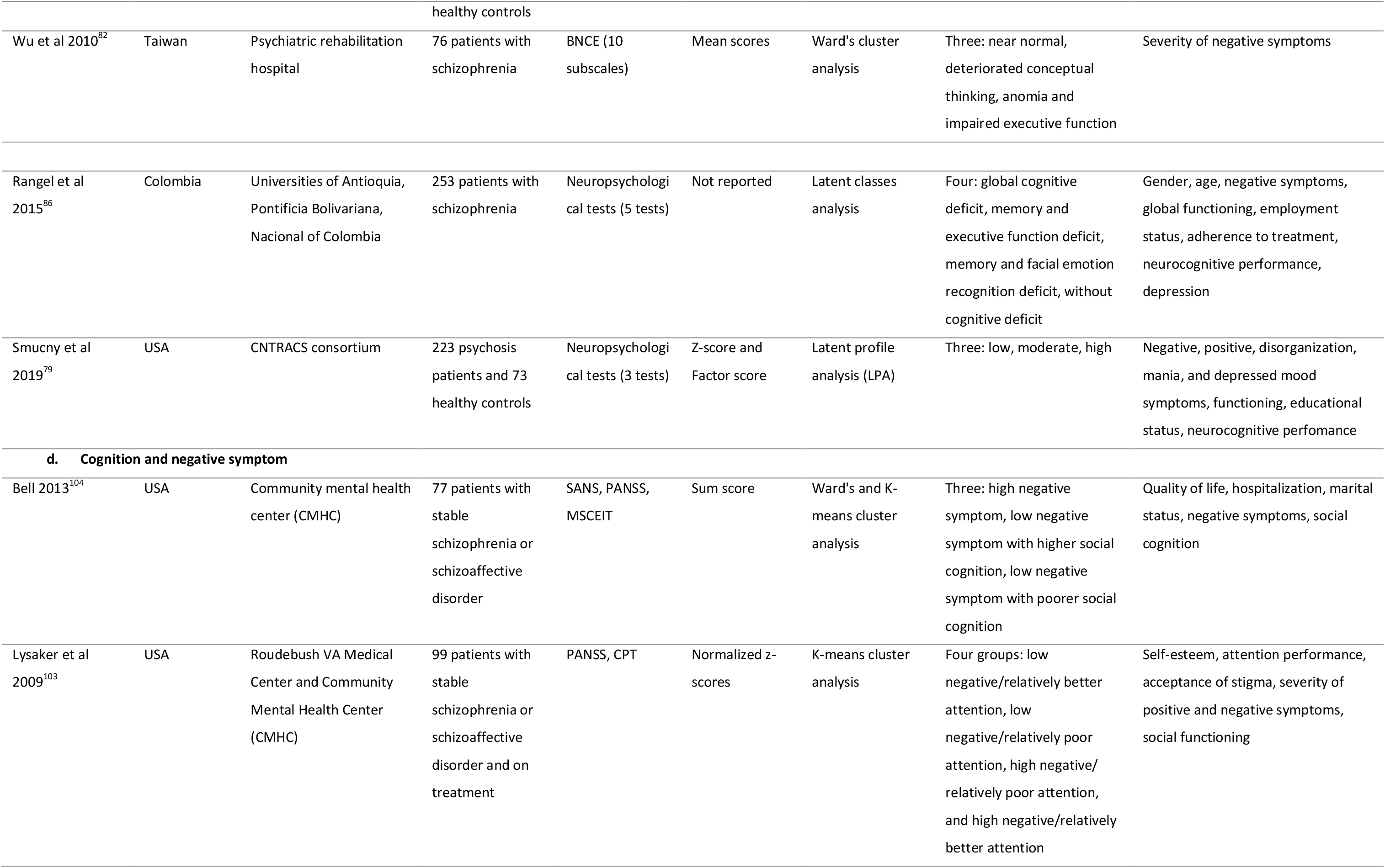
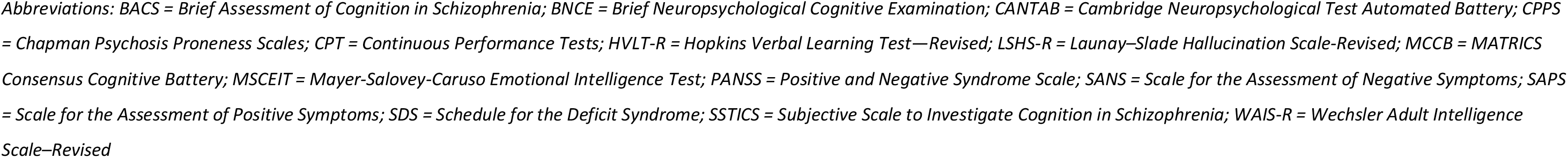
Detailed characteristics of cross-sectional studies (n = 33).

The reported clustering methods were K-means or non-hierarchical clustering analysis^35, 65, 67, 88, 90, 93, 95, 100–103^, Ward’s method or hierarchical analysis^82, 83, 87, 89, 92, 98, 99^, K-means clustering and Ward’s method^32, 37, 66, 80, 85, 91, 94, 97, 104^, latent class or profile analysis^29, 79, 86^ and two-step cluster analysis.^84, 96^ One study^81^ identified clusters using a combination of clinical/empirical and clustering methods. The model selection criteria or similarity metrics were visual inspection of dendrogram, Pearson correlation, squared Euclidean distance, agglomeration coefficients, Dunn index, Silhouette width, Duda and Hart index, elbow test, variance explained, inverse scree plot, average proportion of non-overlap, Akaike information criterion (AIC), Bayesian information criterion (BIC), sample size adjusted Bayesian (ABIC), Schwarz’s Bayesian information criterion (BIC), Lo–Mendell–Rubin (LMR) test, adjusted LMR and the bootstrap likelihood ratio test (BLRT). Squared Euclidean distance was the most common index used to determine the number of clusters.

Of these 21 studies on neurocognitive deficits, 16 studies^37, 65, 79–84, 87–90, 92, 95, 96^ found three clusters, five studies^32, 85, 86, 91, 93^ reported four clusters and one study^94^ discovered five clusters of patients. One study found three clusters in unaffected siblings based on neurocognitive function.^66^ Two studies^29, 97^ reported three clusters of patients based on the negative symptom dimension. Regarding positive symptoms, only one study^98^ identified three clusters of patients and two clusters in the general population. One study^104^ found three clusters of patients by combining social cognition and negative symptom whereas another study^103^ found four clusters of patients based on neurocognition and negative symptom. In addition, two studies^35, 99^ reported three clusters while another study^100^ found out four clusters by combining both positive and negative symptoms. Moreover, three studies^67, 101, 102^ consistently reported four clusters of unaffected siblings or general population based on positive and negative schizotypy dimensions. Generally, the identified clusters had low, mixed (intermediate) and high symptom profiles. Details has been presented in Table 2.

### Predictors of schizophrenia symptoms subgroups

#### Predictors of symptomatic trajectories

Based on evidence from longitudinal studies (Figure 3)^31, 33, 34, 36, 68–77^, the most common identified predictors of severe positive and/or negative symptoms trajectories were older age, male gender, ethnic minority, late age of illness onset, diagnosis of schizophrenia, long duration of untreated psychosis, long duration of illness, poor premorbid, global functioning, and quality of life, low cognitive performance, and severe baseline positive and negative symptoms. Furthermore, gender was identified as a predictor of positive and negative schizotypy in one study.^78^ Regarding neurocognitive impairment, patients with poor trajectories had younger age, low educational status, non-Caucasian ethnicity, lived in a sheltered facility, low IQ, poor premorbid adjustment, severe positive and negative symptoms, and low baseline neurocognitive performance.^30, 64^ Likewise, siblings with poor neurocognitive trajectories had younger age, female gender, low educational status, non-Caucasian ethnicity, low IQ, poor premorbid adjustment, severe schizotypy, frequent positive psychotic experience, and low baseline neurocognitive performance.^64^

**Figure 3:**
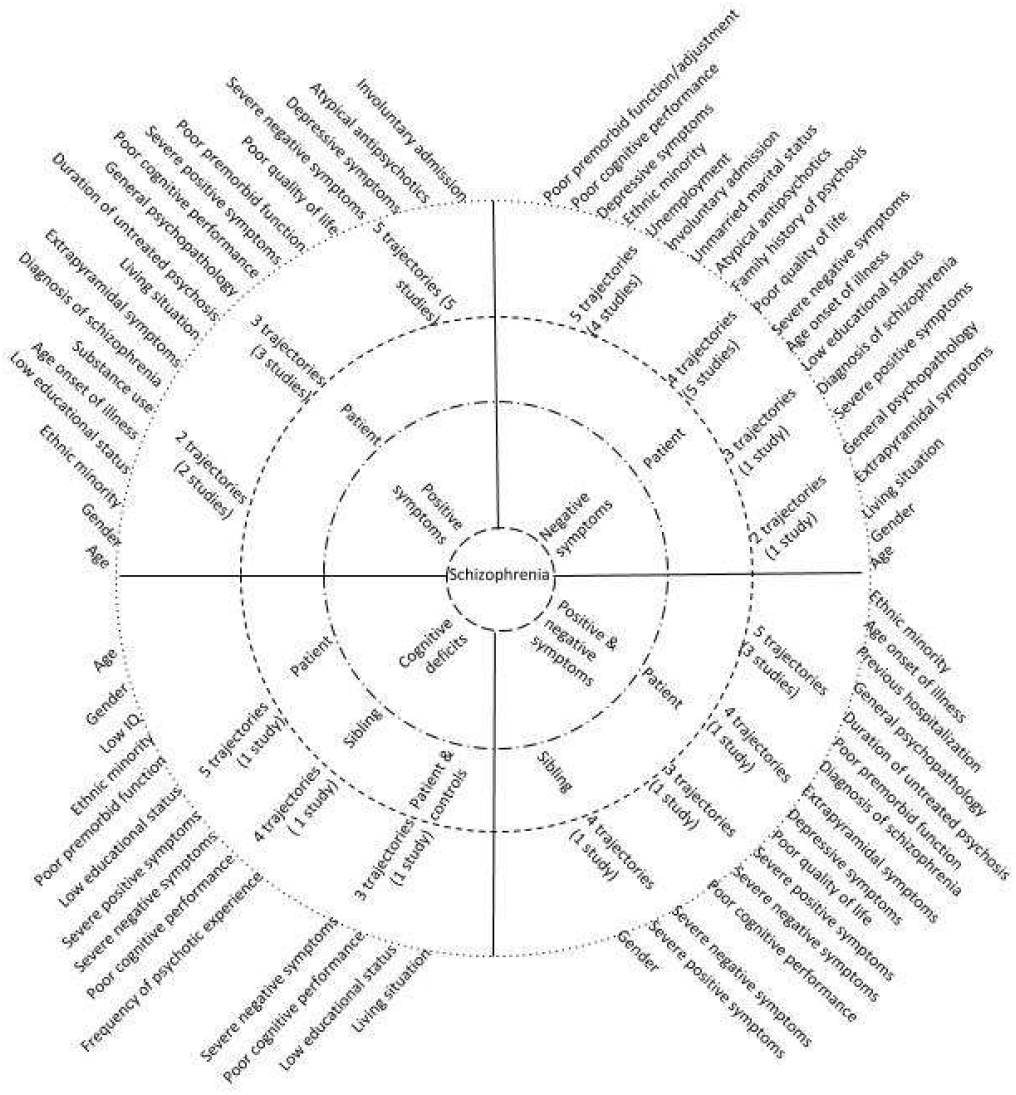
Schizophrenia spectrum circle illustrating predictors of symptomatic trajectories.

#### Predictors of symptomatic clusters

As illustrated in Figure 4, severe positive and/or negative symptoms cluster(s) were predicted by male gender, ethnic minority, low educational status, early age onset of illness, low IQ, severe general psychopathology,, and poor cognition, premorbid adjustment and global functioning.^29, 35, 97, 99, 100^ Severe positive and/or negative schizotypy cluster(s) in unaffected first degree relatives of patients with schizophrenia were predicted by poor experience of pleasure and emotional expression, and low neurocognitive performance.^67^ In the non-clinical population, severe positive and/or negative schizotypy cluster(s) were predicted by male gender, severe paranoid and schizoid symptoms, major depressive episode, substance abuse, medication use, poor social adjustment, severe somatic and anxiety symptoms, and poor neurocognitive and social functioning.^101, 102^

**Figure 4:**
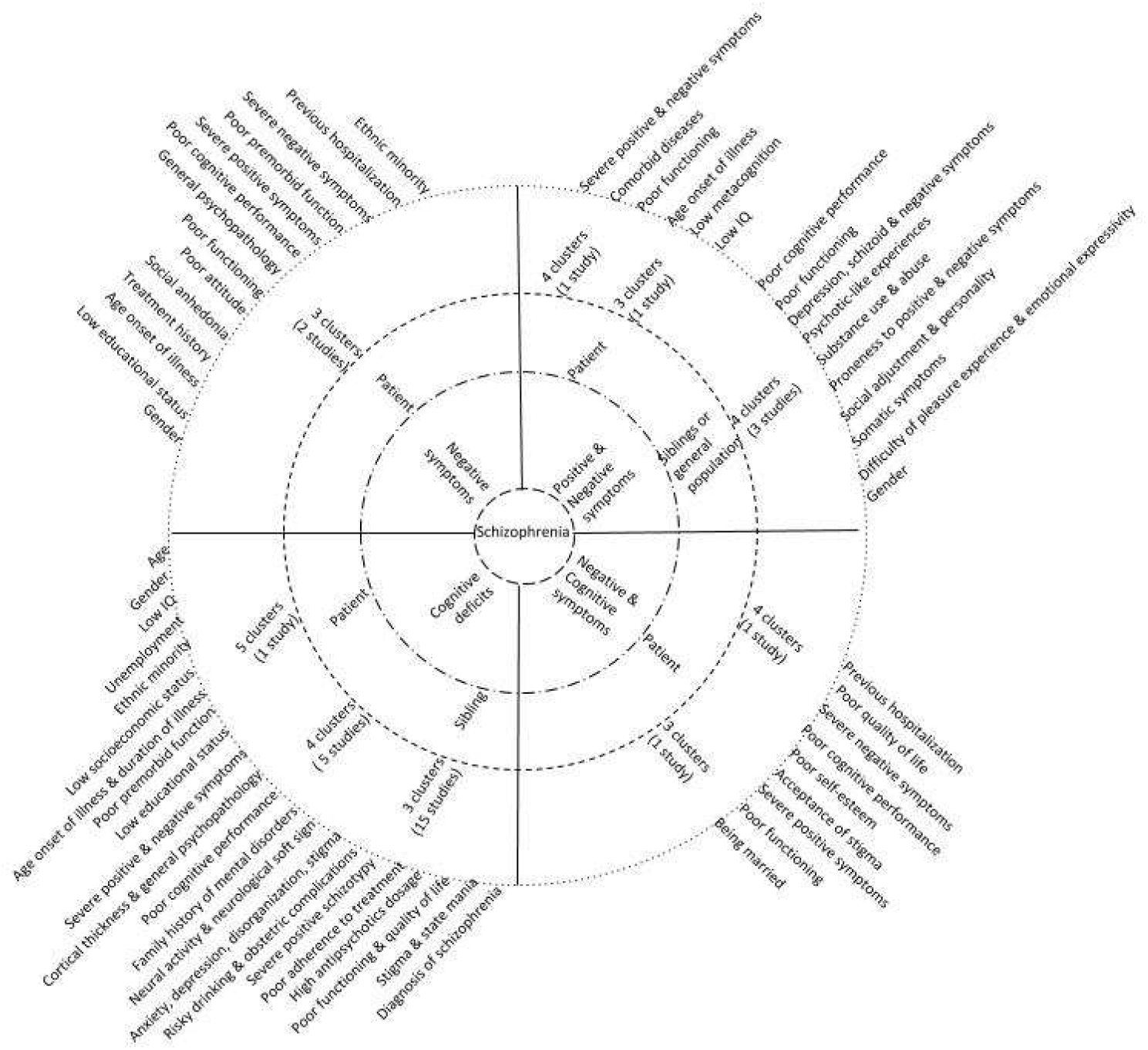
Schizophrenia spectrum circle illustrating predictors of symptomatic clusters.

In addition, poor cognitive impairment cluster(s) were predicted by age, gender, non-Caucasian ethnicity, low socioeconomic and educational status, poor premorbid adjustment, low premorbid and current IQ, early age of illness onset, long duration of illness, severe positive and negative symptoms, poor social cognition, high antipsychotics dosage, use of second generation antipsychotics, and poor functioning and poor quality of life.^32, 37, 65, 66, 79–96^ Siblings subgroups with impaired neurocognitive function were predicted by young age, low educational status, low IQ, poor premorbid adjustment, and severe positive schizotypy (Figure 4).^66^

Overall, as shown in Table 3, 57 predictors of clusters or trajectories were identified by longitudinal and cross-sectional studies across all study participants and symptom dimensions. The most common predictors were old age, male gender, non-Caucasian ethnicity, low educational status, late age of illness onset, diagnosis of schizophrenia, several general psychopathology and depressive symptoms, severe positive and negative symptoms, low cognitive performance, and poor premorbid functioning, quality of life and global functioning.

**Table 3:**
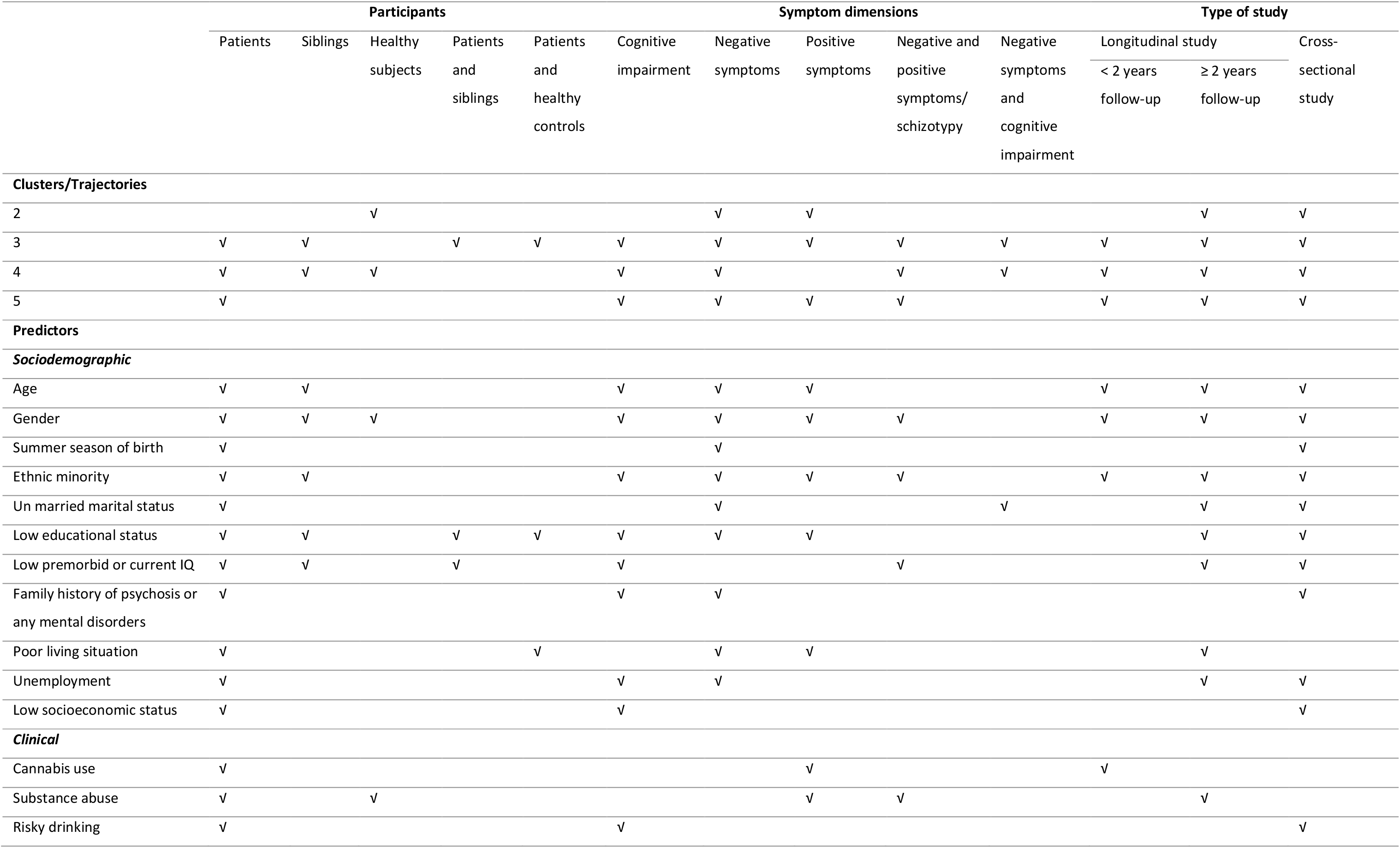
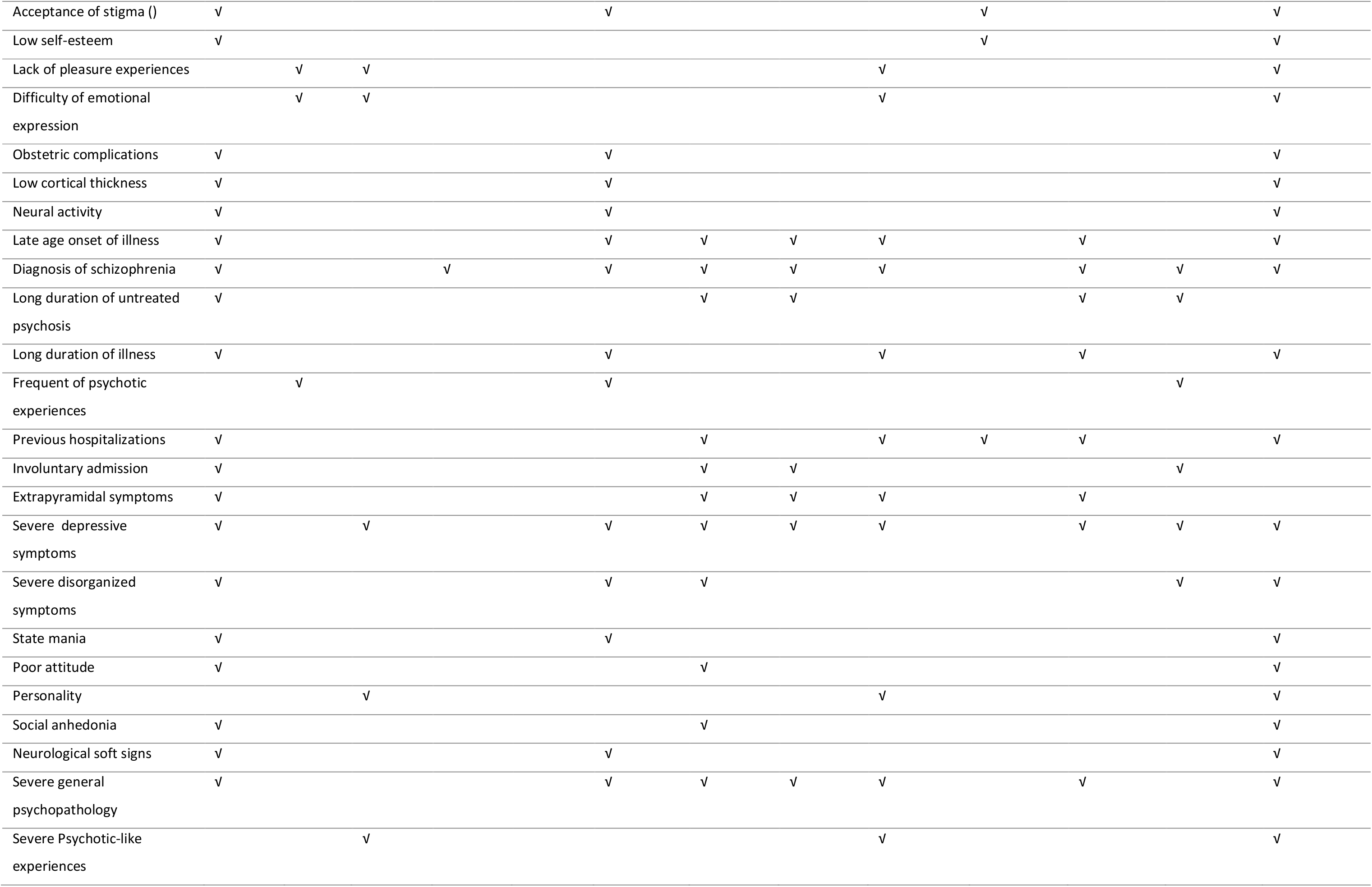
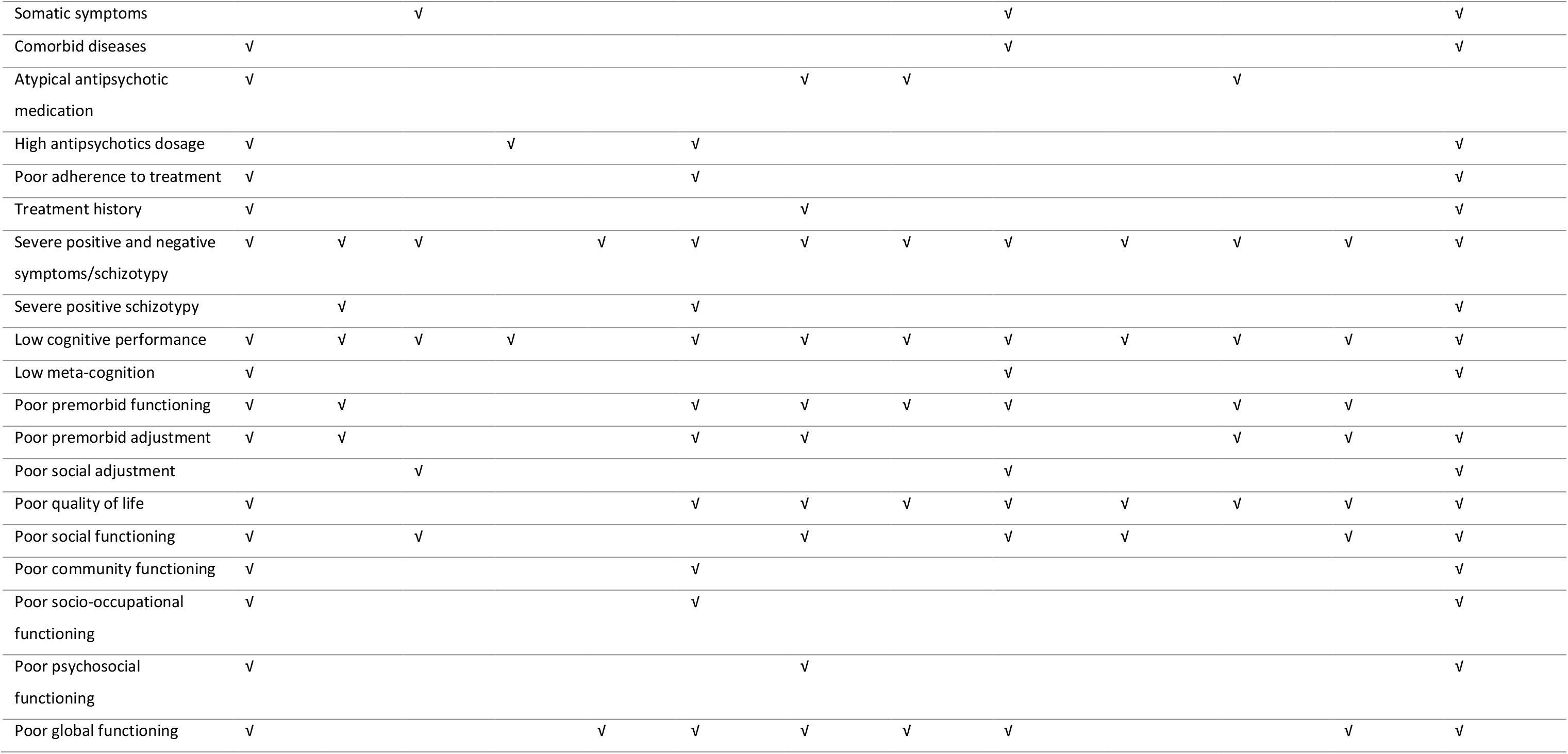
Summary of clusters and trajectories and predictors.

## Discussion

To our knowledge, this is the first comprehensive systematic review on recent cluster- and trajectory-based studies of positive symptoms, negative symptoms and cognitive deficits in patients with schizophrenia spectrum disorders, their siblings and healthy people. Our review has three key findings. First, longitudinal trajectory-based studies distinguished two to five trajectory groups in patients based on positive and negative symptoms, and four to five trajectory groups in patients and siblings based on cognitive deficits. Second, cross-sectional cluster-based studies discovered three clusters of patients based on positive and negative symptoms, and four clusters of siblings based on positive and negative schizotypy. In addition, three to five clusters of patients and their unaffected siblings were discovered based on cognitive deficits. Third, poor symptomatic-outcome trajectories and clusters were predicted by numerous sociodemographic and clinical factors.

We showed that longitudinal studies with patients and siblings have inconsistently identified two to five trajectories across the schizophrenia symptoms. Several shortcomings may cause this inconsistency. Only one-third of the reviewed studies were longitudinal and only two studies^30, 64^ investigated the trajectories of cognitive deficits. This paucity of longitudinal studies on cognitive function may be caused by the fact that neuropsychological assessment is resource intensive, time-consuming, requires specialized data collection training and commitment by study participants. For example, some studies^37, 85, 93^ administered up to 18 psychometric tests, which took more than four hours per wave of assessment. Utterly, none of the reviewed longitudinal studies validated their model against empirical methods or comparable statistical method, and used complex trajectory modelling analysis. Our review showed that growth mixture modelling (GMM)^31, 69, 74^, latent class growth analysis (LCGA)^30, 33, 34, 70, 73, 76, 77^, mixed mode latent class regression modelling^36, 68, 72^ and group-based trajectory modelling (GBTM) were applied.^64, 71, 75^ The difference in patient characteristics may also affect the number of clusters. For example, a studies that included only first-episode psychosis or chronic patients may identify smaller clusters than studies that included a mixture of patients with first-episode and chronic psychosis. Moreover, the difference in frequency and duration of follow-up may lead to subtle difference in results.

Given the scarcity of longitudinal studies, conducting cross-sectional studies and identifying meaningful clusters is the reasonable alternatives. Cluster analysis, which includes K-means clustering and Ward’s method, is data-driven approach for classifying individuals into homogeneous groups by determining clusters of participants that display less within-cluster variation relative to the between-cluster variation.^89^ K-means cluster analysis is a non-hierarchical form of cluster analysis, which is appropriate if previous evidence or hypotheses exist regarding the number of clusters in a sample. It produces the number of clusters initially called for by minimizing variability within clusters and maximizing variability between clusters.^103^ Ward’s method is a hierarchical cluster analysis aiming to determine group assignment without prior hypothesis.^103^ K-means iterative cluster analyses handle larger data sets better than Ward’s method.^102^ To this end, even though they do not to show variability over time, cross-sectional studies are capable of unraveling the heterogeneity of schizophrenia symptoms if appropriate statistical procedures are followed. To date, 33 cross-sectional studies were conducted that found three to five clusters in patients and four in siblings across schizophrenia symptoms. Cognitive deficit was the most commonly examined symptom dimension in cross-sectional studies, whereby 26 studies identified clusters used either K-means^35, 65, 67, 88, 90, 93, 95, 100–103^ or Ward’s method clustering analysis.^82, 83, 87, 89, 92, 98^ Nine cross-sectional studies^32, 37, 66, 80, 85, 91, 94, 97, 104^ cross-validated their model using K-means and Ward’s clustering analysis. Another study^81^ used a combination of clustering and clinical experience to identify homogeneous subgroups.

Longitudinal and cross-sectional studies consistently found several predictors of poor symptomatic trajectories or clusters among patients, unaffected siblings, and general population, including age, gender, ethnic minority, low educational status, late age of illness onset, diagnosis of schizophrenia, severe general psychopathology and depressive symptoms, severe positive and negative schizotypy/symptoms, low cognitive performance, and poor premorbid functioning, quality of life and global functioning. These factors may be used to develop risk prediction model for clinical practice and study disease pathway.

We showed that previous studies included various groups of study population, such as patients with first-episode psychosis or chronic schizophrenia, antipsychotic naïve patients or patients who were on antipsychotic treatment for a month or longer, patients from different age groups and ethnicities, and healthy siblings and controls. While the comparison of patient clusters and trajectories with healthy siblings or controls could provide an accurate means of disentangling the heterogeneity and causes of heterogeneity of schizophrenia symptoms, only four studies (three were cross-sectional studies) examined clusters in siblings. Likewise, most studies used healthy controls to standardize patients neurocognitive composite scores, and few other studies used controls to compare the distribution of patient clusters or trajectory groups. Substantial differences between studies were also noted in constructing composite scores, use of model selection criteria and method of parameter estimation. Moreover, we observed several ways of subtyping and nomenclature for clusters or trajectories, which may be difficult for clinicians to translate the evidence in diagnosing and treating diseases. This is due to the lack of standardized reporting procedures for data analysis plans or results.^54^

The results of statistical subtyping approaches, such as cluster or trajectory analysis depend on mathematical assumptions, type of data, number of variables or tests, sample size and sampling characteristics. Therefore, the models can be unstable and parameter estimates of clinical symptoms may not converge to a consistent set of subgroups and lack a direct relationship to clinical reality.^73, 91, 105^ For example, intermediate clusters and trajectories substantially vary between studies.^91^ We advocate that study results should be applicable, comparable, generalizable and interpretable into clinical practice. We also propose to validate models using additional comparable statistical methods, combine statistical methods of subtyping with empirical methods, and work together with clinicians to create a common understanding and clinically relevant clustering or trajectories nomenclature. Furthermore, it is relevant to replicate clusters or trajectory groups using independent samples, different assessment tools that measure the same construct and different linkage methods.^37, 106^ Finally, further studies are required that focus on longitudinal study design, unaffected siblings and genetic markers as a predictor.

## Conclusions

Our study reveals that schizophrenia symptoms are more heterogeneous than currently recognized and clinically divergent. Future clinical approaches may benefit from the subgrouping of patients to implement person-based therapy. Uncovering the biological basis of individual symptoms may be more helpful in understanding the pathophysiology of the illness than forcing a constellation of co-occurring symptoms.^1^ The identified predictors could be used for developing clinical risk prediction and network modelling, deep endophenotyping, and machine learning to understand symptom pathways. This study showed evidence for clinicians to optimize the efficacy of personalized psychiatric care by predicting individual susceptibility to disease, providing accurate assessment, initiating early intervention strategies, and selecting treatments targeting subgroups of patients with similar phenotypic or psychosocial characteristics.^107^ Therefore, using clustering and trajectory analysis methods will help in implication of precision medicine, in treating subgroups of patients with poor outcome and diagnosing prodromal symptoms in their relatives. Finally, given that personalized psychiatry is at the infancy stage, findings from our review could assist in informing personalized and preventive strategies for clinical practice.^1, 108^

## Funding

Tesfa Dejenie was supported by the Scholarship of University of Groningen, Groningen, the Netherlands. L.H. Rodijk was supported by the Junior Scientific Master Class of the University of Groningen, Groningen, the Netherlands.

## Acknowledgments

We would like to forward our special gratitude to Sjoukje van der Werf, who is medical information specialist at the University of Groningen, the Netherlands, for her support to develop the search strings and guiding the overall literature retrieval process. The authors have declared that there are no conflicts of interest.

## Notes

#### Summary of Updates

This version included updated information and illustrative figures.

https://www.crd.york.ac.uk/prospero/display_record.php?ID=CRD42018093566

